# High throughput isolation of male gametophyte cells of *Solanum lycopersicum* var. Micro-Tom by fluorescence-activated cell sorting

**DOI:** 10.1101/2025.02.12.637952

**Authors:** María Flores-Tornero, Helena Sapeta, Maria Lobato-Gómez, Beatriz Teixeira, Marta Monteiro, Antonio Granell, Jörg D. Becker

**Author notes:** corresponding authors Jörg D. Becker, María Flores-Tornero.

## Abstract

Efficient isolation of male gametes has enabled unprecedented advances in omics research, crucial for elucidating the molecular mechanisms governing male gametogenesis and fertilization. In this study, we developed a method for isolating generative and sperm cells from the economically important crop *Solanum lycopersicum*.

A double fluorescent marker line was generated in the tomato variety Micro-Tom, employing mTurquoise and mScarlet-I fluorescent proteins under the control of promoters exhibiting preferential activity in generative and sperm cells, respectively. Then we developed a protocol that combines male gamete release from pollen of this line and SYTOX Red live/dead cell stain to obtain viable cells by Fluorescence-activated cell sorting. This allows the isolation of generative cells from mature pollen grains, and of sperm cells from pollen tubes after semi-*in vivo* growth, both in high quantity and purity. Additionally, an unexpected mScarlet-I signal in the vegetative nucleus, that persists until the sperm cells are formed, allows the sorting of vegetative nuclei.

We anticipate that our novel double-marker line will accelerate research into tomato male gametogenesis, thereby enhancing efforts to improve the resilience of fertilization processes to climate change.

## INTRODUCTION

Male gametophyte development consists of microsporogenesis, where a microsporocyte undergoes meiosis to form four microspores, and subsequent microgametogenesis, involving two pollen mitoses (Boavida, Becker, and Feijó 2005). The first mitosis produces a generative cell (GC), and a vegetative nucleus, essential to create a pollen tube to transport the male gametes to the ovule. During the second mitosis, the GC divides and produces two sperm cells (SC) that later on will fuse with the female gametes and finalize the double fertilization process. In some species, like *Arabidopsis thaliana*, these two mitoses are already performed in the mature pollen before anthesis, resulting in tricellular pollen grains consisting of a vegetative nucleus and the two SC. However, other species, like *Solanum lycopersicum*, produce bicellular pollen containing only the GC and the vegetative nucleus (Lu, Wei, and Wang 2015). In those species, the second mitosis occurs at some point during the progamic phase, a series of events starting with pollen germination on the stigma, pollen tube growth through the pistil, and SC delivery inside the ovule. In most cases, this second mitosis can be observed in pollen germinated *in vitro* using specific germination media, whereas in others the pollen requires the female stimulus, germinating on the top of a cut style and allowed to grow through the female tissue to trigger the second mitosis in semi-*in vivo* conditions (Cao, Reece, and Russell 1996).

The continued improvements of omic analysis allow cell profiling at an unprecedented level of detail, improving our understanding of cell-specific omic changes and molecular mechanisms regulating development and function under specific conditions. In the case of male reproductive cells, traditional difficulties in isolating them and the low amount recovered were the main problems to characterize these specific cells (Flores-Tornero and Becker 2023). Among cell isolation methods, micromanipulation is one of the most popular ones because it is inexpensive and allows high purity samples of single cells, but it is time consuming and yields a low number of cells. Another popular method is based on Percoll gradient and has been established in tomato to isolate high amounts of male gametes (Lu, Wei, and Wang 2015). However, the similar morphology of SC and GC and the lack of available markers made very difficult to guarantee a high purity of those isolations. Recently, fluorescence-activated cell sorting (FACS) has become an interesting option among the tricellular pollen species, because it enables specific isolation of high amounts of male gametes using fluorescent dyes (Engel et al. 2003) or cell-specific fluorescent marker lines (Borges et al. 2012). In the case of Arabidopsis, this method allowed not only the characterization of the SC transcriptome but also the spliceosome, highlighting the specificity of the molecular regulation in these cells (Misra et al. 2019, 2022; Borges et al. 2008). Generating such information for crop species like tomato is crucial for uncovering the molecular regulation of male reproductive cells development, and fertilization mechanisms such as sperm-egg recognition and fusion.

Research on model plants like Arabidopsis already benefit from having well characterised genomes, availability of cell-specific fluorescent marker lines, and detailed protocols to isolate male gametes and study the fertilisation process at microscopic and molecular level. However, relevant crops like tomato are lacking some of these powerful tools, limiting our understanding of the fertilisation process, and the possibility of manipulating it in our favour. Here we report the creation of the first sperm, generative cell, and vegetative nucleus fluorescent marker line in tomato (Micro-Tom variety) and our protocol for high throughput isolation by FACS. With this contribution we aim to facilitate future omic profiling of male reproductive cells and the exploration of the fertilisation process in tomato.

## MATERIAL AND METHODS

### Plant material and growth conditions

Micro-Tom tomato lines were sown on soil (SIRO professional multiplication mixture: Humus, Peat, Perlite,) and grown in controlled conditions, with a temperature of 25/20ºC (day/night) and 60% relative humidity. A 14-hour light period (14h/10h day night) was used with 200 µmol/m2/s average light intensity at plant level. After 8 weeks, plants started to flower, and pollen/flowers were used during the flowering period (additional 6 weeks). Watering was performed three times a week with tap water.

### Cloning and plant transformation

All the primers and plasmids described here are listed in **Supplementary Table 1**. 1,000bp of the promoter of the selected genes *Solyc07g008580*.*3*.*1 (GCp)* and *Solyc01g112000*.*4*.*1 (SCp)*, as well as the coding sequence of mTurquoise fluorescent protein were amplified from genomic DNA using Phusion High-Fidelity DNA Polymerase (NEB) and the primers were designed using the GoldenBraid Pro cloning webpage (https://goldenbraidpro.com/)(SCp_DOMfw/rev, GCp_DOMfw/rev, and mTur_DOMfw/rev, respectively). The amplicons were purified from the gel and cloned separately into pUPD2 vectors, generating the entry clones “pUPD2_SCp”, “pUPD2_GCp”, and “pUPD2_ mTurquoise”, respectively. Then, the transcriptional unit *SCp::Scarlet-I* was created by combining “pUPD2_SCp”, “pUPD2_Scarlet-I”, and “pUPD2_T35S” into the “alpha2” vector via GoldenBraid Pro assembly using BsaI restriction sites. Similarly, the transcriptional unit *GCp::mTurquoise* was created by combining “pUPD2_GCp”, “pUPD2_ mTurquoise”, and “pUPD2_T35S” into an “alpha2” vector.

Next, the “alpha2_ GCp:: mTurquoise” vector was combined with the plasmid “alpha1R_nptII” containing antibiotic resistance into the final vector “omega1” by using BsmbI restriction sites. In parallel, the “alpha2_ SCp::Scarlet-I” was cloned into an “omega 2” vector by using the twister plasmid GB0106 and BsmbI restriction sites, in order to make both constructs compatible in the last destination vector. Finally, the “omega1_ GCp:: mTurquoise_nptII” vector was cloned in tandem with the “omega2_ SCp::Scarlet-I” into the final vector “alpha1”, and sequenced before transforming *Agrobacterium*.

Tomato transformation was carried out according to the protocol described in (Martí et al. 2007). In brief, tomato seeds were germinated *in vitro*, and cotyledons were inoculated with *Agrobacterium tumefaciens* harbouring the construct of interest. Following inoculation, the cotyledons were transferred to selective media for promoting shoot and root development. Putative transgenic plantlets were then genotyped for the presence of the transgenes using the primers listed in **Supplementary Table 1**. Transgenic plantlets rooting in selective media were subsequently transferred to the greenhouse for further experiments.

### Microscopy

Pollen grains and pollen tubes were placed in 35mm imaging Petri dishes (MatTek) filled with pollen germination media (PGM: 0.2 mM Ca(NO_3_)_2_, 0.1 mM KNO_3_, 0.2 mM MgSO_4_, 0.2 mM H_3_BO_3_, 10% sucrose and 1% Casein) and stained with the nucleic acid dye SYBR green. Images of the time course were taken with an inverted fluorescence microscope TE300 (Nikon). Microscopy experiments were performed at the Bioimaging Unit of the Gulbenkian Institute for Molecular Medicine (Oeiras, Portugal). Pollen grains and pollen tubes of the double fluorescent tomato marker line were imaged with a 3i Marianas SDC microscope. They were excited with the 488-nm, 561-nm and 445-nm lasers, and the emitted light was measured at 497-522 nm for SYBR Green, 570-593 nm for mScarlet-I and 457-502-nm for mTurquoise, respectively.

### Generative cell and sperm cell isolation

GC were isolated from mature pollen grains obtained from fully open flowers at anthesis, using the double marker line. Pollen from 3 to 4 flowers was collected by vibration (**Fig.1C**) and resuspended in sperm cell buffer (1.3 mM H_3_BO_3_, 3.6 mM CaCl_2_·2H_2_O, 0.74 mM KH_2_PO_4_, 438 mM Sucrose, 7 mM MOPS, 0.83 mM MgSO_4_·7H_2_O, pH adjusted to 6.0) (Santos, Bispo, and Becker 2017). Then, the pollen wall was broken mechanically by continuous vortex at 2200 rpm for 2 min with glass beads (425–600μm, Sigma), according to Santos et al. (2017). Afterwards, the solution was filtered with a 15 µm filter, placed on ice and samples were used for GC sorting.

SC from the double marker line were isolated from semi-*in vivo* grown pollen tubes. Closed flower buds were emasculated 72 hours prior to anthesis. Emasculated flowers at the anthesis stage were pollinated with freshly collected pollen and placed on double side tape. Pistils were excised at the junction of the style and ovary and placed on PGM for 17.5 hours. Afterwards pollen germination medium was replaced by burst solution (0.45M mannitol supplemented with 0.5% cellulase, 0.3% macerozyme R-10, and 0.05% pectolyase Y-23 enzymes) and incubated for 30 min at 25°C. Then the suspension was transferred to 400 µl of ice-cold SC extraction buffer and filtered through a 15 µm filter.

A detailed step by step protocol for generative cell and sperm cell isolation is presented in **Supplementary File 1**.

### Fluorescence-activated cell sorting

Single-cell suspensions of GC from mature pollen grains or SC from pollen tubes prepared as previously described were stained with 5 nM SYTOX Red (Invitrogen) for 10 minutes on ice. Cells were sorted in a 5-laser Cytek Aurora CS (Cytek Biosciences). The spectral signature of the unstained sample was assessed using unlabelled cells of the WT tomato plants isolated with the exact same methodology, which were also used to prepare the single-stained control for the viability dye. The single stained controls for mScarlet-I and mTurquoise were prepared, respectively, with SC or GC isolated from the single marker lines. The gain settings for the physical parameters were defined using the mScarlet-I labelled cells, back-gating the mScarlet-positive events in the YG2-A detector onto the FSC-A vs. SSC-A dot plot. For the fluorescence detectors, we used the Cytek Assay Settings. To isolate mScarlet-I cells, events with an intermediate side scatter profile and a high fluorescence intensity in the YG2-A detector, which collects the mScarlet-I peak emission, were gated; these events were subsequently analysed in a dot plot displaying the R1-A vs. SSC-A detectors, selecting those with low fluorescence intensity in R1, which correspond to the cells that did not incorporate the dead-cell exclusion reagent; finally, we excluded groups of two cells together (doublets) by displaying the selected events in a SSC-A vs. SSC-H dot plot and used this final gate for sorting. Isolation of the mTurquoise labelled cells followed a different gating strategy. First, we identified GC based on FSC-A and SSC-A; second, we excluded dead cells, which display a high fluorescence intensity in the R1-A detector and an intermediate SSC profile; third, we selected the events with high fluorescence intensity in the V4-A detector, which collects the mTurquoise peak emission; and finally, we excluded doublets as described above. The presence of mScarlet-I or mTurquoise in the sorted cells was validated by sorting samples directly into 35 mm microscopy plates (MatTek) filled with sperm cell buffer and immediately observed under a 3i Marianas Lightsheet microscope (settings details in the microscopy section).

## RESULTS

### GC divide into two SC 14 hours after germination

In *Solanum lycopersicum* var. Micro-Tom **(Fig. 1A)**, hundreds of pollen grains are formed inside the bilobed anthers that form a cone surrounding the long style in each flower (**Fig. 1B**). After applying a slight vibration on open flowers **(Fig. 1C)**, pollen grains are released on the stigma while many others fall out of the flower and can be harvested for analysis. As tomato produces bicellular pollen, GC and a vegetative nucleus are already present, and can be observed and isolated directly from mature pollen **(Fig. 1D)**. However, SC become visible only after the GC divides inside the pollen tube at some point after germination. Therefore, it is necessary to induce the germination of pollen, and this can be done either by incubating the pollen in a specific germination media (*in vitro*) **(Fig. 1E)**, or by pollinating the stigma of a cut style (semi-*in vivo*) and placing it on germination media to allow pollen tubes to emerge from the cut end **(Fig. 1F; Supplementary File 1)**.

**Fig 1.**
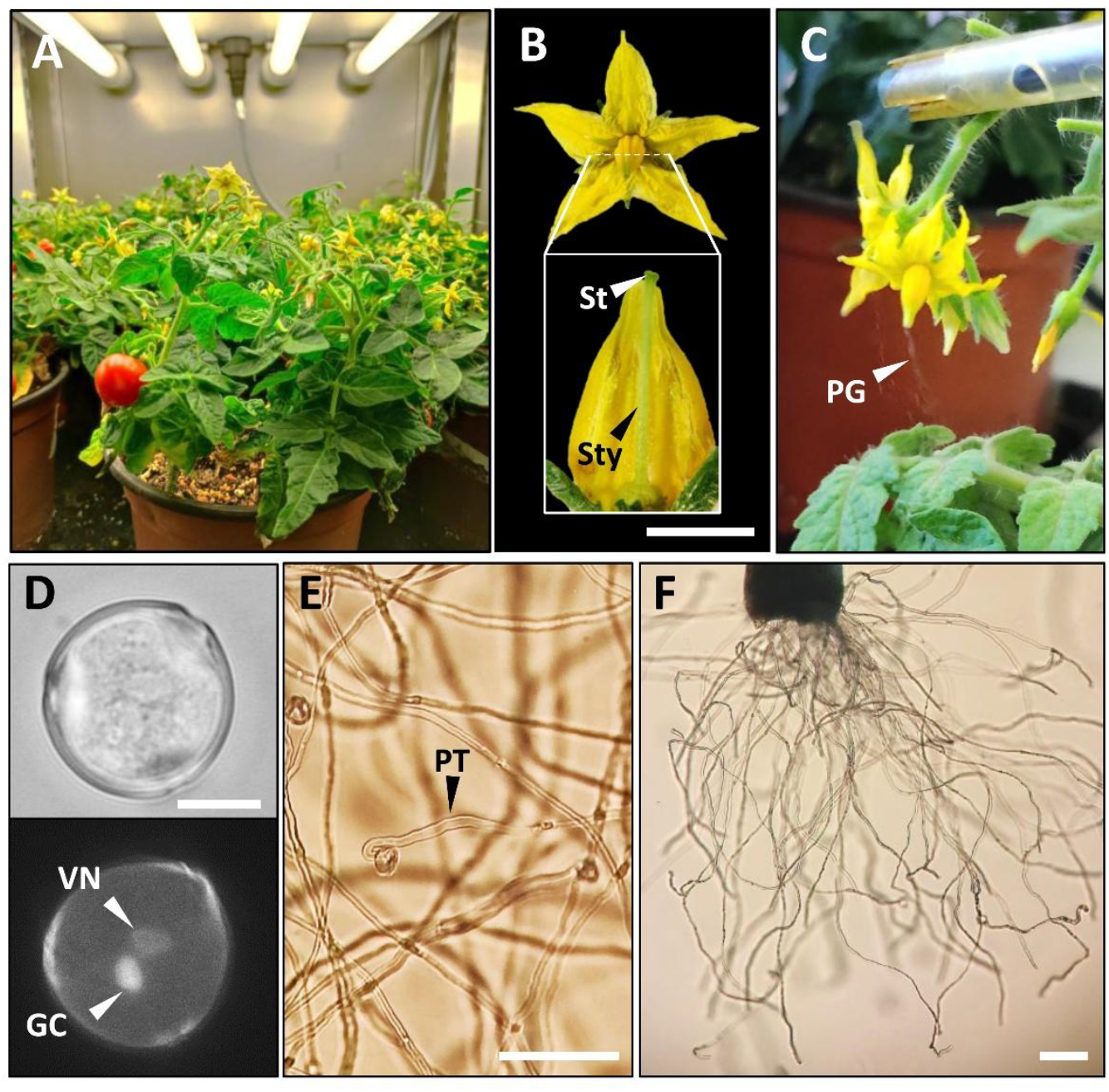
Micro-Tom flower, pollen and pollen tube growth *in vitro* and semi-*in vivo*. A) Micro-Tom plant growing in a reach-in chamber. B) Structure of a mature flower (up), showing the conic anther surrounding the style and the ovary at the base (down). C) Pollen release by the vibration method. D) Bright field (up) and fluorescent field (down) of a bicellular pollen stained with DAPI, showing the generative cell and the vegetative nucleus. E) Micro-Tom pollen tubes growing *in vitro* and F) in semi-*in vivo*. GC, generative cell; PG, pollen grain; PT, pollen tube; St, stigma; Sty, style; VN, vegetative nucleus. Scale bars in B: 75 mm; D: 10 µm; E and F: 100 µm.

Next, we identified the specific time point at which the highest percentage of SC can be observed. We performed a time course germination of pollen grains *in vitro* and semi-*in vivo* and tracked the division of the GC with the nucleic acid stain SYBR Green **(Fig. 2)**. SC started to be observed inside the pollen tubes 14 hours after germination *in vitro* **(Fig. 2A)**, but in a very low proportion (1%, data not shown). However, at this time point in the semi-*in vivo* experiment **(Fig. 2B)**, the proportion of observed SC reached 75% (**Supplementary Table 2**), thus being the preferred germination system to isolate high amounts of SC. Semi-*in vivo* pollen tubes showed 90% presence of SC 18 hours after germination; thus, this time point was selected for sperm cell isolation.

**Fig 2.**
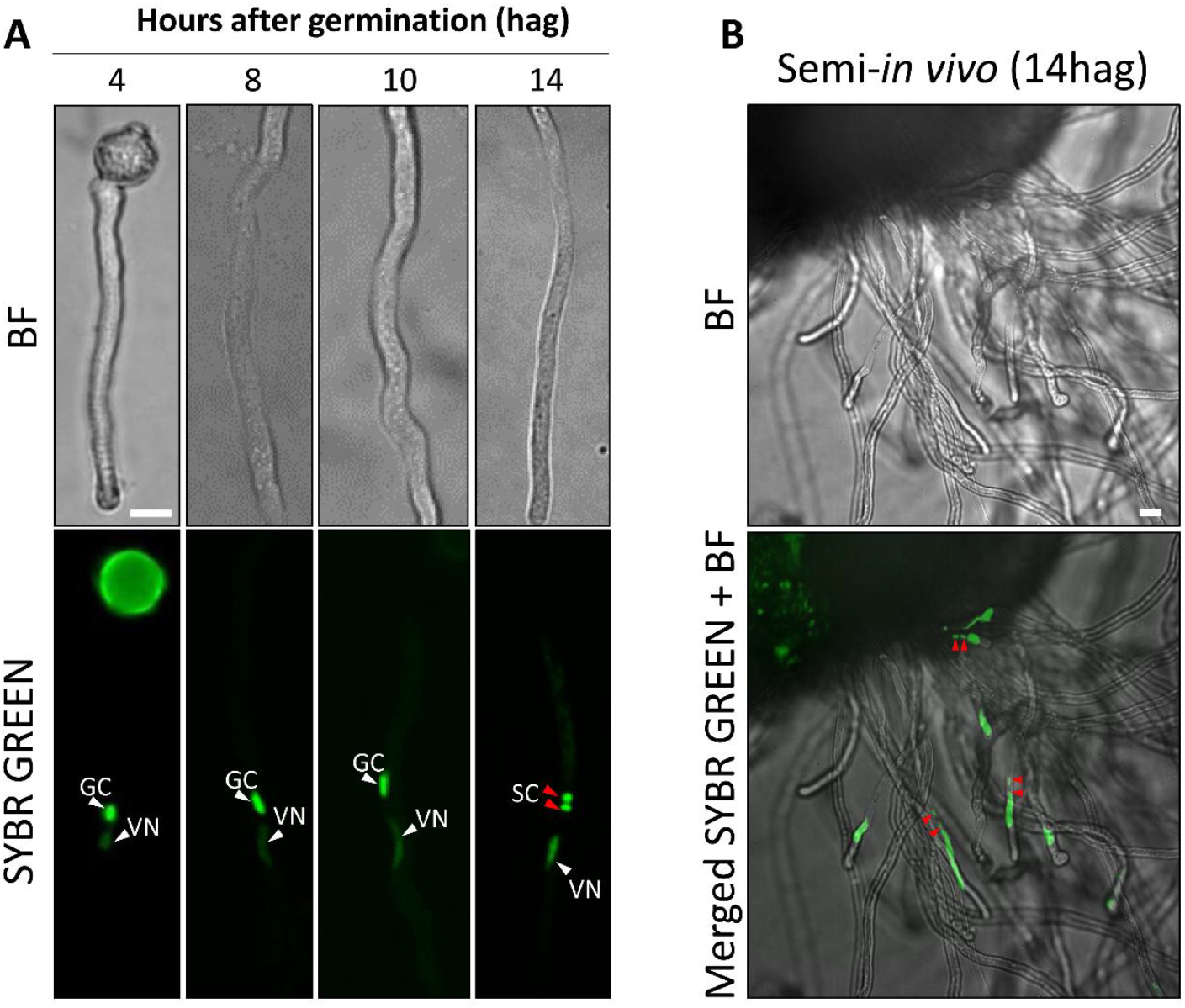
Time course of pollen tube germination *in vitro* and semi-*in vivo*. A) Generative cells divide 14hag, producing the two sperm cells necessary for double fertilisation. B) Semi-*in vivo* assay showing emerging pollen tubes from the cut style and the presence of sperm cells. Scale bar = 10um.

### Fluorescent proteins facilitate the identification of GC and SC

The specific expression of a fluorescent protein considerably improves the identification of a cell type within a mixture of cells. In tomato pollen grains, there is a GC and a vegetative nucleus, and most of the GC will divide later to produce two SC, but many others will not. The result is a mixture of three different types of cells, thus the need for an efficient identification to isolate them.

To differentiate generative and SC, we created a double fluorescent marker line in which the fluorescent proteins mTurquoise and mScarlet-I are under the control of promoters preferentially active in either GC or in SC, respectively. To this end, we used the EvoRepro database (https://evorepro.sbs.ntu.edu.sg/) (Julca et al. 2021) to screen for candidate genes in the transcriptomic profile of tomato generative and SC compared to other tissues (**Supplementary Table 3**). First, we filtered those genes (263) that were preferentially expressed in SC with a specificity measure (SPM) = 0.85 (Xiao et al. 2010). Then, we sorted those genes by high expression, first in SC and then in GC, finally focussing on the top 25 candidates. As there are no genes expressed exclusively in SC, we obtained the expression ratio “sperm cell/generative cell” for each gene and selected those with the highest and the lowest ratio. In this way, we selected *Solyc01g112000*.*4*.*1* (alpha-like-class expansin) because it is 13.23 times more expressed in SC, whereas the gene *Solyc07g008580*.*3*.*1* (class II/ASH1 histone methyltransferase component of histone lysine methylation/demethylation) is 21 times more expressed in GC. By choosing these two extremes, our goal was to maximize the difference between these two cells by using two different fluorescent proteins under two promoters differentially expressed in both cell types.

Then, we used the GoldenBraid GB3.0 approach (Vazquez-Vilar et al. 2017) to clone the promoter of *Solyc01g112000*.*4*.*1*, preferentially expressed in SC *(SCp)* controlling the expression of mScarlet-I, together with the promoter of *Solyc07g008580*.*3*.*1*, preferentially expressed in generative cell *(GCp)*, to control the expression of mTurquoise **(Fig. 3A)**. After tomato transformation, we obtained three independent double marker lines named *gcsc 13, gcsc 17*, and *gcsc 16*, with a strong fluorescent signal in GC **(Fig. 3B)** and SCs **(Fig. 3C)** in more than 95% of the samples. Interestingly, we could also observe a strong fluorescence mScarlet-I signal in the vegetative nucleus in all three independent lines **(Fig. 3B)**, which gets considerably reduced when the SC are formed.

**Fig 3.**
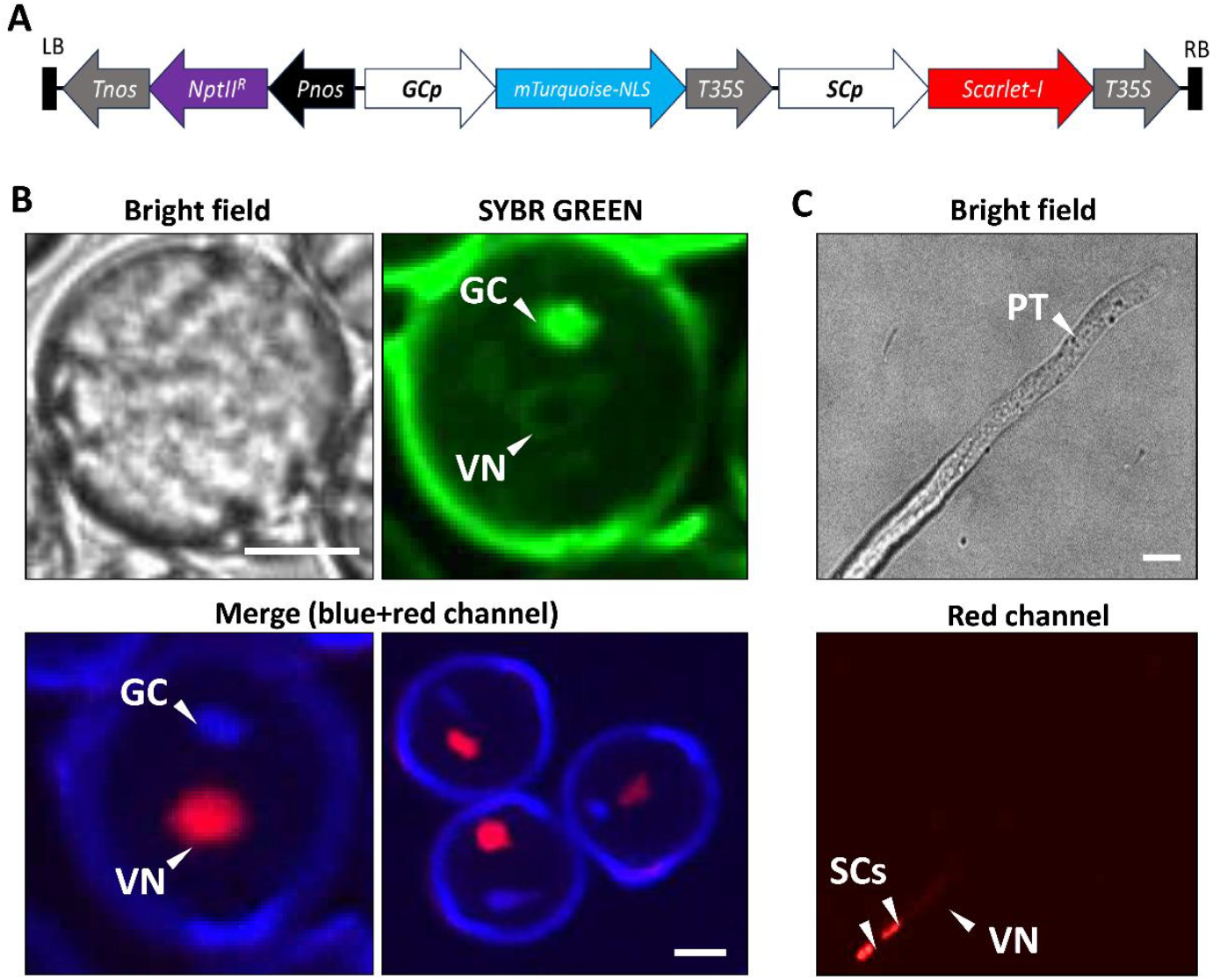
GCSC fluorescent tomato marker line. A) Genetic construct of the double marker line. B) Bicellular pollen grain from the Micro-Tom double marker line GCp::mTurquoise-NLS SCp::Scarlet-I. C) Pollen tube 14hag showing two sperm cells. GC, generative cell; VN, vegetative nucleus; SC, sperm cells; NptIIR, neophosphinotricine; PT, pollen tube. Scale bar (10 µm).

### Optimized method for FACS isolation of male gametes

Viable GC were isolated from mature pollen grains, and SC were isolated from semi-*in vivo* grown pollen tubes using fluorescence-activated cell sorting (FACS). For GC, the gating strategy involved identifying the GC population based on forward (FSC-A) and side scatter (SSC-A). Cell viability was then performed using a live/dead stain (SYTOX Red), with non-viable cells excluded based on their high R1-A detector signal. Afterwards, mTurquoise-expressing GC were distinguished by their higher fluorescence intensity in the Violet 4 (V4-A) detector (**Fig. 4A**). A different gating strategy was used for the isolation of SC. The cells population was identified using FSC-A and SSC-A, and later mScarlet-I-expressing SC were detected due to higher fluorescence intensity in the Yellow Green 2 (YG2-A) detector (**Fig. 4B**). Cell viability was then assessed as described for GC, with non-viable cells excluded based on their high R1-A detector signal. For both cell types, doublets were also excluded, and only viable singlets were sorted (singlets population, **Fig. 4**).

**Fig 4.**
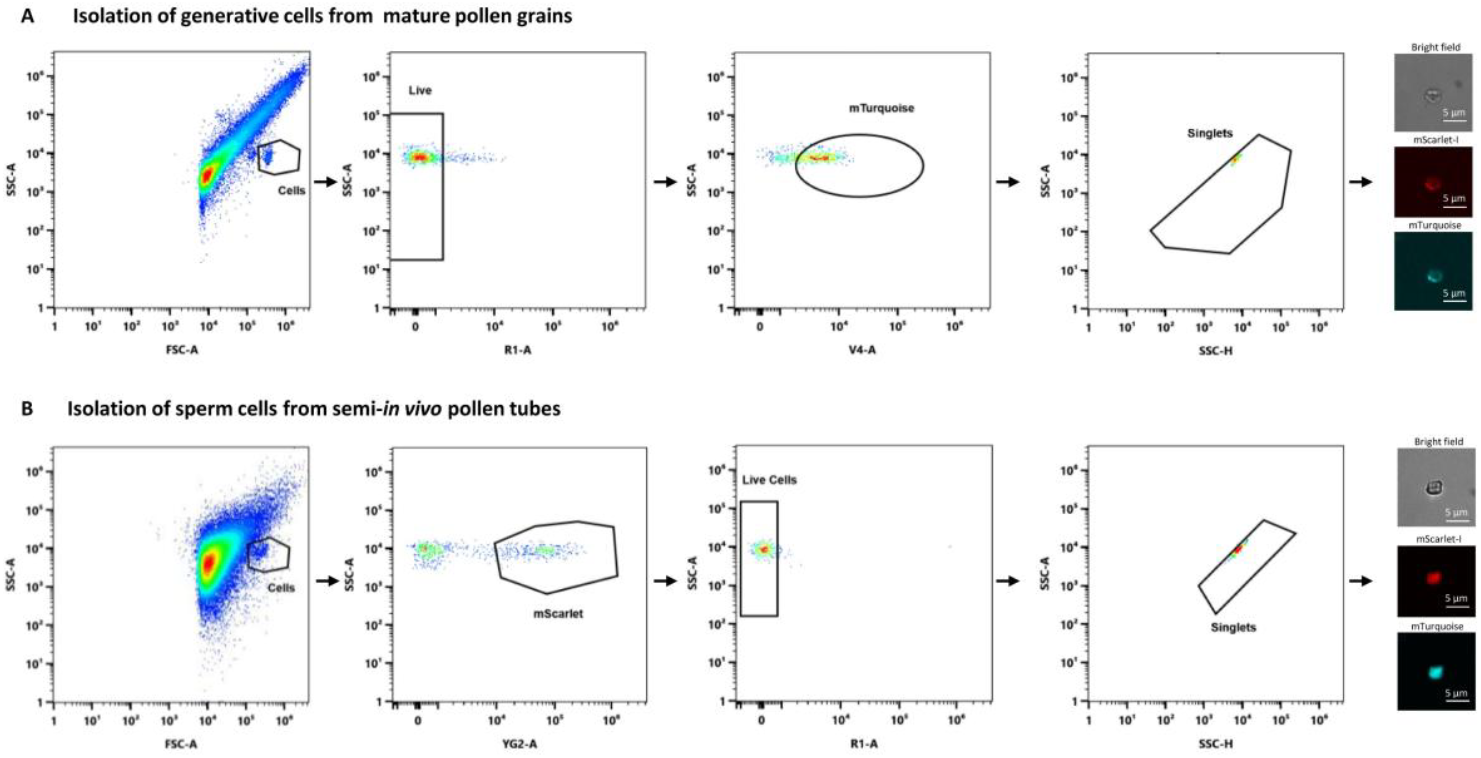
FACS isolation of viable generative cells from mature pollen grains and sperm cells from semi-*in vivo* grown pollen tubes. A) Gating strategy to isolate viable generative cells. Cells were first discriminated from debris based on forward (FSC-A) and side scatter (SSC-A) parameters; then, non-viable cells were excluded based on the incorporation of SYTOX red and consequent fluorescence emission in R1-A detector, allowing selection of live cells (Live Cells population); afterwards, mTurquoise-expressing generative cells were identified based on their high fluorescence intensity in the Violet 4 (V4-A) detector (mTurquoise population); finally, we excluded doublets based on SSC-A and SSC-H, ensuring that only single cells were sorted (Live-singlets population). The identity of sorted cells was confirmed by observation under the microscope. B) Gating strategy to isolate viable sperm cells. First, the cell population was identified and separated from debris based on FSC-A and SSC-A; second, mScarlet-I expressing sperm cells were identified because of higher fluorescence intensity in the Yellow-Green 2 (YG2-A) detector (mScarlet SC population); third, live cells were identified (Live Cells population) and SYTOX red non-viable cells were excluded (R1-A detector signal); finally, doublets were excluded based on SSC-A and SSC-H, and only single cells were sorted (Live-singlets population).

The purity of sorted generative and SC was confirmed by microscopy. We observed mTurquoise and mScarlet-I signal under the microscope for both cell types. This is observed because, as explained above, there are no genes expressed exclusively in generative / SC and the selected promotors were based on the expression ratios between both cell types. Yet when we compare the spectral profile of the generative and SC populations in the cytometer, SC exhibited significantly higher mScarlet-I expression compared to GC, as highlighted by their strong signal in the yellow-green region (YG1 to 9, **Supplementary Figure 1**). These observations confirm the increased expression of mScarlet-I in SC as compared to GC.

GC from mature pollen grains can also be isolated via FACS using nucleic acid dyes such as SYBR Green (data not shown). However, due to the coexistence of generative and SC in pollen tubes, isolating a pure sperm cell population via FACS requires a double marker line.

## DISCUSSION

The goal of exploring the omic profile of GC and SC has been addressed already in some species (Flores-Tornero and Becker 2023), but difficulties in separating those cells and isolating large amounts with less material represented a considerable limitation.

Some studies with pollen of *S. lycopersicum* var. Heinz1706 developed a percoll density gradient method to separate SC, GC, and vegetative nucleus (Lu, Wei, and Wang 2015). However, due to the similar morphology of SC and GC, the purity of the isolations was difficult to ensure. They used pollen tubes 10 hours after germination, as 92% of them contained SC. In our case, Micro-Tom required at least 18 hours after germination to observe similar percentages of SC under semi-*in vivo* germination conditions. It is known that for some species with bicellular pollen, GC division can be triggered *in vitro* by adding reduced forms of nitrogen (Read, Clarke, and Bacic 1993). Moreover, the influence of the female tissue during the semi-*in vivo* assays considerably increased the division of the GC, and in certain species, GC division is entirely contingent upon female influence (Lin et al. 2020).

In the last years, FACS sorting has proven to be an excellent alternative to sort cells dyed with DAPI or Hoechst. However, some concerns arose regarding potential interferences on immune precipitations assays involving chromatin (Mari et al. 2010) as well as unforeseen preferences of DAPI for protein aggregates rather than nucleic acids (Mora et al. 2020), making specific labelling less reliable. Nowadays, available databases are a great source of information to pick candidates and generate gamete marker lines to sort this type of cells. In this regard, the use of cell-specific promoters to express fluorescent proteins to label a cell type unambiguously bypasses the use of dyes and any potential negative effect in downstream omic analysis.

In principle, the aim of this study was to create a double marker line to allow the differentiation between GC and SC and improve their sorting efficiency. Surprisingly, we also observed a strong red fluorescence signal in the vegetative nucleus in three independent lines, supporting it is a *bona fide* signal and discarding a positional effect. The sperm cell specificity of the promoter selected for this study and its none-specificity in vegetative nucleus is further confirmed by transcriptomic data from other tomato studies (Song et al. 2024). This result can indicate a potential exchange of material from the vegetative nucleus to SC, a theory that has been already suggested (McCue et al. 2011; Jiang et al. 2015).

In summary, this double marker line enables the sorting of not only GC and SC but also vegetative nuclei of *S. lycopersicum* var. Micro-Tom pollen, e.g. to study the negative effect of climate change in the fertilization process by omics analyses.

## Abbreviations

FACS: Fluorescence-Activated Cell Sorting
GC: generative cell
SC: sperm cells

## ACKNOWLEDGEMENTS

We acknowledge the technical support of Patricia Rodrigues from the Bioimaging Unit of the Gulbenkian Institute for Molecular Medicine, funded by PPBI-POCI-01-0145-FEDER-022122. We also thank the Bacterial Imaging Cluster (BIC) hosted at the Instituto de Tecnologia Química e Biológica António Xavier (ITQB-NOVA), especially to Carolina Feliciano for microscopy assistance. This work was partially supported by PPBI - Portuguese Platform of BioImaging co-funded by national funds from OE - “Orçamento de Estado” and by European funds from FEDER - “Fundo Europeu de Desenvolvimento Regional”. This work was supported by the Fundação para a Ciência e a Tecnologia (FCT), I.P., project grants PTDC-ASP-PLA-2007-2020, and CEECIND/03345/2018 to JDB, GREEN-IT - Bioresources for Sustainability R&D Unit base (DOI: 10.54499/UIDB/04551/2020) and programmatic (DOI: 10.54499/UIDP/04551/2020) funding, LS4FUTURE Associated Laboratory (DOI: 10.54499/LA/P/0087/2020), as well as by the European Union’s HORIZON-MSCA-2021-PF-01-01 programme (grant 101059247 to MF-T). We acknowledge support through the EU-funded project HARNESSTOM (101000716), Innovation Action EC-H2020-SFS-2020-1, and the Ministerio de Ciencia e Innovación (Spain) through Agencia Estatal de Investigación (PID2022-141438OB-I00 BTC). M.L.G. and A.G. are participants of the European COST action CA18210 (ROXY) for networking activities. M.L.G is recipient of an ACIF-2021 fellowship (GVA, Generalitat Valenciana).

## AUTHOR CONTRIBUTIONS

JDB and MFT conceived the study. MFT performed the cloning, microscopy and wrote the manuscript together with HS, with input of the other authors. HS selected, grew and propagated the fluorescent marker lines. BT, MM and HS developed the FACS isolation method. MLG and AG did the Micro-Tom transformations.

## CONFLICT OF INTEREST

We declare no conflict of interest

## DATA AVAILABILITY

All data are available in the main text and in the supplementary material. Request for seeds should be done to Jörg D. Becker.

## Notes

### Competing Interest Statement

The authors have declared no competing interest.

