## Supplementary Figure 1 for "High throughput isolation of male gametophyte cells of *Solanum lycopersicum* var. Micro-Tom by fluorescence-activated cell sorting"

A Spectrum profile of generative cells preferentially expressing mTurquoise

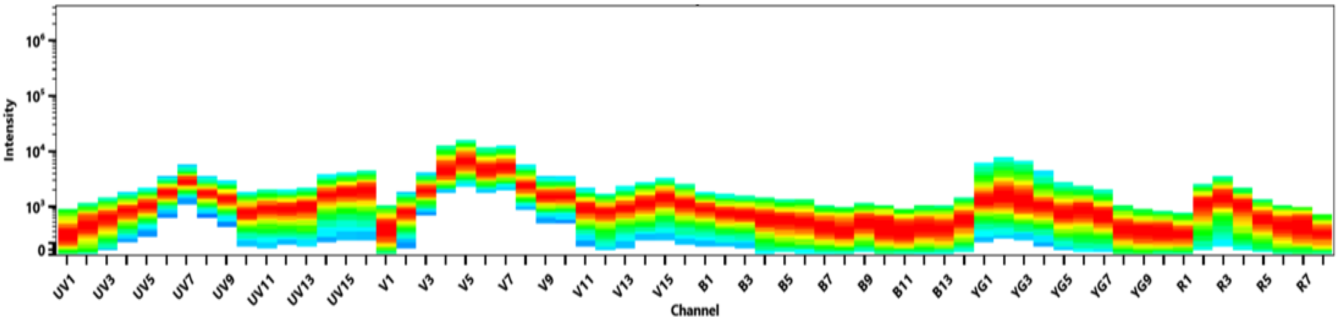


B Spectrum profile of sperm cells preferentially expressing mScarlet-I

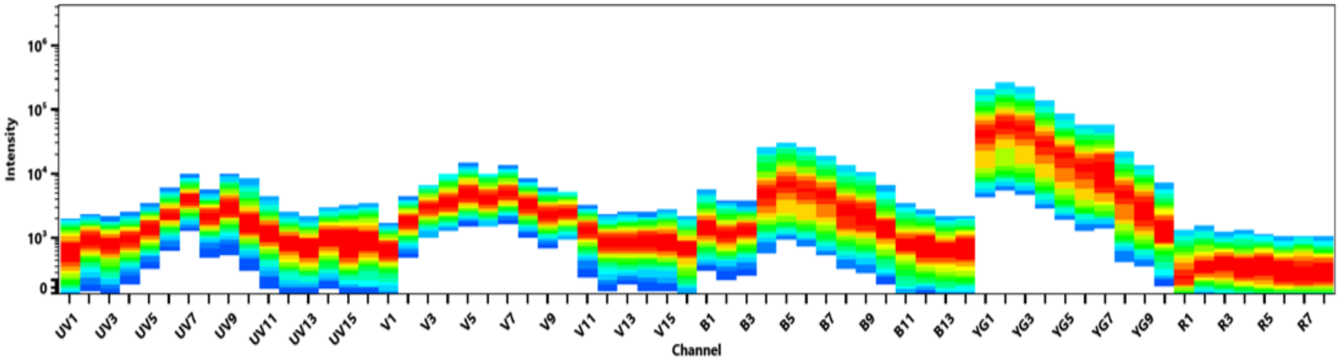


**Supplemental Figure 2:** Spectral profiles obtained using the 5-laser Cytek Aurora CS (Cytek Biosciences) for the mTurquoise (A) and mScarlet-I (B) populations (refer to Figure 4). The profiles correspond to generative cells isolated from pollen predominantly expressing mTurquoise and sperm cells isolated from pollen tubes predominantly expressing mScarlet-I, respectively.
