## Supplementary Table 1 for "High throughput isolation of male gametophyte cells of *Solanum lycopersicum* var. Micro-Tom by fluorescence-activated cell sorting"

**Supplementary Table 1. List of primers and plasmids used in this study**

| Oligo name | Sequence (5' -> 3') |
|---|---|
| SCp_DOMfw | GCGCCGTCTCGCTCGGGAGCCCGCACACGGAATAACAGA |
| SCp_DOMrev | GCGCCGTCTCGCTCACATTATTTATTTATTTTTTGTTATATGAAGAAATGAATT |
| GCp_DOMfw | GCGCCGTCTCGCTCGGGAGTCCCTTTTTACTTGTCCACTTTTG |
| GCp_DOMrev | GCGCCGTCTCGCTCACATTCTATGCAACCTTTTGCCAGAA |
| mTur_DOMfw | GCGCCGTCTCGCTCGAATGGCGTCCAAAGGGGAAGA |
| mTur_DOMrev | GCGCCGTCTCGCTCAAAGCTCATTATTCTTCTTCTTGGTCAGC |
| pUPD2_FW | GCTTTCGCTAAGGATGATTTCTGG |
| pUPD2_Rev | CAGGGTGGTGACACCTTGCC |
| LB_omega1 | GGTGGCAGGATATATTGTGG |
| SCpmid_FW | CCGTGTATTGAGCATTACATTCC |
| Scar_1 | AAGAAGCCCGTGCAGATG |
| nptII_Rev | ACCTGCCCATTCGACCACCAA |
| RB_Rev | GTTTACCCGCCAATATATCCTGT |
| GCpend_FW | GGTTGAACGTAATCATACTTTTCT |
| mTur_mid_FW | GGCGGACCATTATCAACAGAA |
| SC-Scarletti_FW | CCGTTCAGTGGCACCCATTA |
| SC-Scarletti_Rev | TGTAGTCCTCGTTGTGGGAG |
| GC-mTur_FW | CATCTGCGCATAGCGAAAGC |
| GC-mTur_Rev | CATCTGCGCATAGCGAAAGC |

| Plasmid ID* | Plasmid name | Description |
|---|---|---|
| GB0015 | Alpha1 | |
| GB0017 | Alpha2 | |
| GB0019 | Omega1 | |
| GB0021 | Omega2 | |
| GB0036 | pUPD2_T35S | |
| GB0106 | Twister alpha2 to omega | |
| GB0107 | Twister from alpha1 to omega | |
| GB0158 | Twister from omega2 to alpha | |
| GB0226 | Alpha1_nptii | |
| GB0466 | Twister from omega1 to alpha | |
| GB4048 | pUPD2_Scarletti | |
| GB0307 | pUPD2 | Domesticator plasmid with chloramphenicol resistance. |

*All this plasmids and additional information can be found in the online database GoldenBraid Pro (https://goldenbraidpro.com/).
