## Supplementary Table 2 for "High throughput isolation of male gametophyte cells of *Solanum lycopersicum* var. Micro-Tom by fluorescence-activated cell sorting"

**Supplementary Table 2**. Time course indicating the percentage of sperm cells in semi-*in vivo* germinated pollen tubes. Cells were counted in tubes grown in at least 6 independent pistils. GC, generative cell; SC, sperm cell.

| Time after germination (h) | Total number of tubes analysed | GC observed (%) | SC observed (%) |
|---|---|---|---|
| 14 | 91 | 25.3 | 74.7 |
| 16 | 172 | 19.8 | 80.2 |
| 18 | 288 | 10.4 | 89.6 |
