## Supplementary File 1 for "High throughput isolation of male gametophyte cells of *Solanum lycopersicum* var. Micro-Tom by fluorescence-activated cell sorting"

**Detailed protocol: Isolation of Generative Cells from mature pollen grains**

1. **Pollen collection**
   - Collect mature pollen grains on a glass petri dish by gently vibrating 3–4 flowers at anthesis.
   - Vibration to release pollen can be achieved by gently touching the flower peduncle with a vibrating object (e.g. a TissueRuptor from Qiagen, or an electrical toothbrush). Alternatively, use small forceps to remove the anthers and tap them into a glass Petri dish to release the pollen.
2. **Add glass beads**
   - Transfer the collected pollen to a microcentrifuge tube and add approximately 150 µl of glass beads (425–600 µm, Sigma).
3. **Add buffer**
   - Add 500 µl of ice-cold sperm cell buffer (Santos et al., 2017) containing:
     - 1.3 mM $H_3BO_3$
     - 3.6 mM $CaCl_2 \cdot 2H_2O$
     - 0.74 mM $KH_2PO_4$
     - 438 mM Sucrose
     - 7 mM MOPS
     - 0.83 mM $MgSO_4 \cdot 7H_2O$
     - Adjust pH to 6.0
4. **Disrupt pollen wall**
   - Vortex the tube at 2200 rpm for 2 minutes to break the pollen wall and release the pollen content, including generative cells.
5. **Filter and isolate Generative Cells**
   - Filter the supernatant through a 15 µm mesh using a CellTrics® filtering system into a 2 ml microcentrifuge tube.
   - Centrifuge briefly (5 seconds) to pass the solution through the mesh to exclude debris.
6. **Place the Generative Cell suspension on ice. The sample is ready for FACS.**

**Detailed protocol: Isolation of Sperm Cells from semi-*in vivo* grown pollen tubes**

1. **Emasculation**
   - Emasculate closed flower buds 72 hours before anthesis by carefully removing the conic anthers with fine forceps (see Detailed Protocol Figure 1A).
2. **Pollen collection**
   - After 72 hours, collect fresh pollen from 8 to 10 flowers as described above and store it in a glass Petri dish.
3. **Collect emasculated pistils**
   - Remove the sepals and collect emasculated pistils by holding them at the stalk.
   - Place the pistils on a glass slide with the ovary fixed on the edge of the double-sided tape (see Detailed Protocol Figure 1B).
4. **Inspect pistils**
   - Using a stereomicroscope, check the pistils for damage and confirm that the stigma is receptive (appears glossy). Discard any damaged or non-receptive pistils.
5. **Pollination**
   - Dip the receptive stigmas into the freshly collected pollen to pollinate them (see Detailed Protocol Figure 1C).
6. **Excise pollinated pistils**
   - Fix the pollinated pistils on the glass slide with double-sided tape.
   - Excise the pistils at the junction of the style and ovary with a sharp needle (the cut must be fast and clean to avoid squashing the style base).
7. **Semi-*in vivo* assembly**
   - Gently place the excised pistils along the edge of a 35 mm imaging Petri dish (MatTek) containing 40 µl of freshly pre-prepared pollen germination medium:
     - 0.2 mM $Ca(NO_3)_2$
     - 0.1 mM $KNO_3$
     - 0.2 mM $MgSO_4$
     - 0.2 mM $H_3BO_3$
     - 10% Sucrose
     - 1% Casein
   - Ensure the cut end of the style is in direct contact with the medium (see Detailed Protocol Figure 1D).

8. **Incubation**
    - Cover the Petri dish with its lid, lined with wet paper to maintain humidity.
    - Place the dish in a humid chamber and incubate at 25°C for 17.5 hours.
9. **Pollen tube burst and cell release**
    - After incubation, remove the pollen germination medium by pipetting.
    - Replace it with 40 µl of burst solution containing:
        - 0.45 M Mannitol
        - 0.5% Cellulase
        - 0.3% Macerozyme R-10
        - 0.05% Pectolyase Y-23
    - Incubate the dish at 25°C for 30 minutes.
    - Pipette the solution up and down gently to ensure the collection of most of the sperm cells.
11. **Collect cell suspension**
    - Transfer the released cell suspension into a collection tube containing a 15 µm mesh using a CellTrics® filtering system.
    - Optionally, repeat the mixing step with an additional 50 µl of sperm cell buffer to maximize recovery of sperm cells.
12. **Add extraction buffer**
    - Add 400 µl of ice-cold sperm extraction buffer to the collected suspension.
13. **Filter and isolate Sperm Cells**
    - Filter the suspension through a 15 µm mesh by brief centrifugation (5 seconds) to exclude debris.
14. **Place the Sperm Cell suspension on ice. The sample is ready for FACS.**

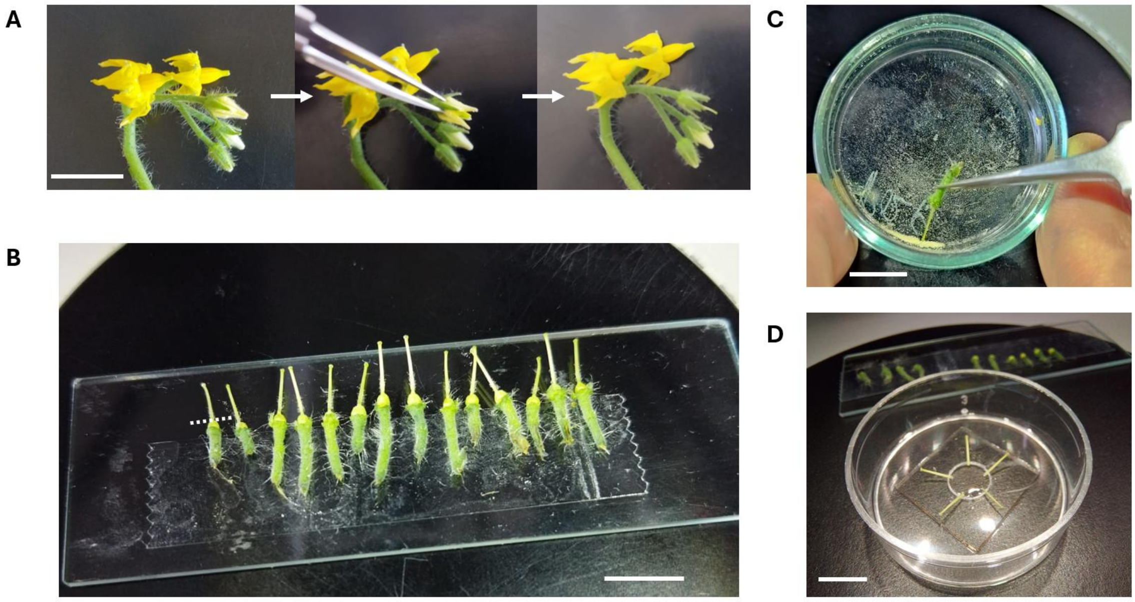


**Detailed Protocol Figure 1: Preparation of semi-*in vivo* grown pollen tubes for sperm cell isolation.** A) Flower emasculation: the conic anthers are removed with a fine forceps from closed flower buds 72 hours before anthesis; B) Collection of emasculated pistils: at the anthesis day, emasculated pistils are collected and placed by the stalk on double side tape; C) Pollination: the pistil stigma is dipped in pollen and placed back on the double side tape. The style is then cut at the ovary base (dashed line panel B); D) Placement of pollinated pistil in medium: cut pollinated pistils are gently placed inside the germination medium with the style base in direct contact with the medium. Scale bar is 1 cm.
